# A Development-Inspired Niche for Homeostatic Human Mini-Intestines

**DOI:** 10.1101/2022.06.12.495827

**Authors:** Charlie J. Childs, Emily M. Holloway, Caden W. Sweet, Yu-Hwai Tsai, Angeline Wu, Joshua H. Wu, Oscar Pellón Cardenas, Meghan M. Capeling, Madeline Eiken, Rachel Zwick, Brisa Palikuqi, Coralie Trentesaux, Charles Zhang, Ian Glass, Claudia Loebel, Qianhui Yu, J. Gray Camp, Jonathan Z. Sexton, Ophir Klein, Michael P. Verzi, Jason R. Spence

**Affiliations:** Department of Cell and Developmental Biology, University of Michigan Medical School, Ann Arbor, MI 48109, USA; Department of Computational Medicine and Bioinformatics, University of Michigan, Ann Arbor, MI, 48109, USA; Department of Internal Medicine, Gastroenterology, University of Michigan Medical School, Ann Arbor, MI 48109, USA; Rutgers, The State University of New Jersey, Piscataway, New Jersey, USA; Department of Biomedical Engineering, University of Michigan Medical School and University of Michigan College of Engineering, Ann Arbor, MI, 48109, USA; Program in Craniofacial Biology and Department of Orofacial Sciences, University of California, San Francisco, CA, USA; Department of Medicinal Chemistry, College of Pharmacy, University of Michigan, Ann Arbor, MI, 48109, USA; Department of Pediatrics, Genetic Medicine, University of Washington, Seattle, WA 98195, USA; Department of Materials Science and Engineering, University of Michigan College of Engineering, Ann Arbor, MI, 48109, USA; Roche Institute for Translational Bioengineering (ITB), Roche Pharma Research and Early Development, Roche Innovation Center, Basel, Switzerland; Institute of Molecular and Clinical Ophthalmology Basel, 4031 Basel, Switzerland

## Abstract

Epithelial organoids derived from intestinal tissue, also referred to as mini-intestines or mini-guts, recapitulate many aspects of the organ *in vitro* and can be used for biological discovery, personalized medicine, and drug development. Murine intestinal organoids represent a homeostatic system that balances stem cell maintenance within a crypt-like compartment and differentiation within a villus-like compartment^1–3^. However, this homeostatic balance and spatial organization has not been achieved with human intestinal organoids^4^. Here, we leverage single cell RNA-seq data (scRNA-seq) and high-resolution imaging to interrogate the developing human intestinal stem cell niche. We identified an EGF-family member, EPIREGULIN (EREG), as uniquely expressed in the developing crypt, and found that EREG can take the place of EGF as an *in vitro* niche factor. Unlike EGF, which leads to growth of thin-walled cystic organoids, EREG-organoids are spatially resolved into budded and proliferative crypt domains and a differentiated villus-like central lumen. Transcriptomics and epigenomics showed that EREG-organoids are globally similar to the native intestine while EGF-organoids have an altered chromatin landscape, downregulate the master intestinal transcription factor *CDX2*^5,6^, and ectopically express stomach genes.

## Body

The intestinal epithelium is organized into crypts, where intestinal stem cells (ISCs) are located, and villi, which contain differentiated cell types that carry out absorptive, secretory, and immunologic functions required for life. ISCs are maintained in an environment known as the stem cell niche, which is a collective of biochemical and physical cues that regulate maintenance, self-renewal, and differentiation^7^. Niche cues include extracellular matrix (ECM), cell-cell contacts, growth factors, cytokines, metabolites, and environmental cues such as food and bacterial products^8^. Understanding stem cell regulation and the stem cell niche has been a focus in the murine intestine^9–12^, and recent work have begun to interrogate the adult and developing human stem cell niche using single cell genomic approaches^13–20^.

Three dimensional epithelial organoids, often referred to as enteroids, mini-guts, or mini-intestines^2,21–23^, have become a powerful *in vitro* tool to interrogate human stem cell biology and the stem cell niche. First described in mice, epithelial organoids feature crypt-like domains that possess stem and Paneth cells budding from a central lumen that contains differentiated cell types (i.e. goblet, enteroendocrine, enterocytes) in a spatial pattern that recapitulates the native tissue^1–3,24–30^. Interestingly, while many studies have described the growth/culture of adult and fetal human epithelial organoids, a similar spatially organized homeostatic state has not been achieved. Instead, human organoids typically grow as thin-walled cysts replete with proliferative stem and progenitor cells but lacking differentiated cell types and spatial organization^4^. Although prior work has identified niche cues that enhance human cell differentiation *in vitro*, the spatial organization observed in murine organoids has not been achived^31,32^.

Here we identify the EGF family member, EPIREGULIN (EREG), as a growth factor uniquely expressed in the stem cell domain of the developing human intestine. By replacing EGF ligand, which is commonly used in organoid culture medium, with EREG, we observed that organoids grow in spatially organized crypt-like and villus-like domains with the full complement of epithelial cell types found in the native intestine. We compare organoids grown in EGF and EREG using single cell transcriptomic and epigenomic approaches and find that EGF-grown organoids have reduced expression of the master intestinal transcription factor, *CDX2*, which is associated with an altered chromatin landscape and ectopic expression of stomach-associated genes. On the other hand, EREG maintained chromatin architecture and gene expression similar to native tissue. Collectively, our data suggest that accurately mimicking the *in vivo* niche leads to more faithful mini-intestines *in vitro* and shows the importance of EGF-family members in human stem cell homeostasis.

### Identification of EREG as a Novel Niche Cue

To better understand the human intestinal stem cell niche, we leveraged single cell RNA-seq data (scRNA-seq) of the developing human intestine^19,33^ and performed Louvain clustering and visualization using uniform manifold approximation and projection (UMAP)^34^. Major cell classes were annotated using cohorts of genes associated with neurons, immune, endothelial, mesenchymal, and epithelial cells (Figures S1d-f). Within the epithelial compartment, we identified ISCs (*LGR5/OLFM4)*, enterocytes (*FABP2/ALPI/RBP2)*, BEST4+ enterocytes (*BEST4/SPIB*), goblet cells (*MUC2/SPDEF/DLL1)*, tuft cells (*TRPM5/TAS1R3*), and enteroendocrine cells (*CHGA/NEUROD1/PAX6/ARX/REG4)* (Figure 1e and S1f).

**Figure 1.**
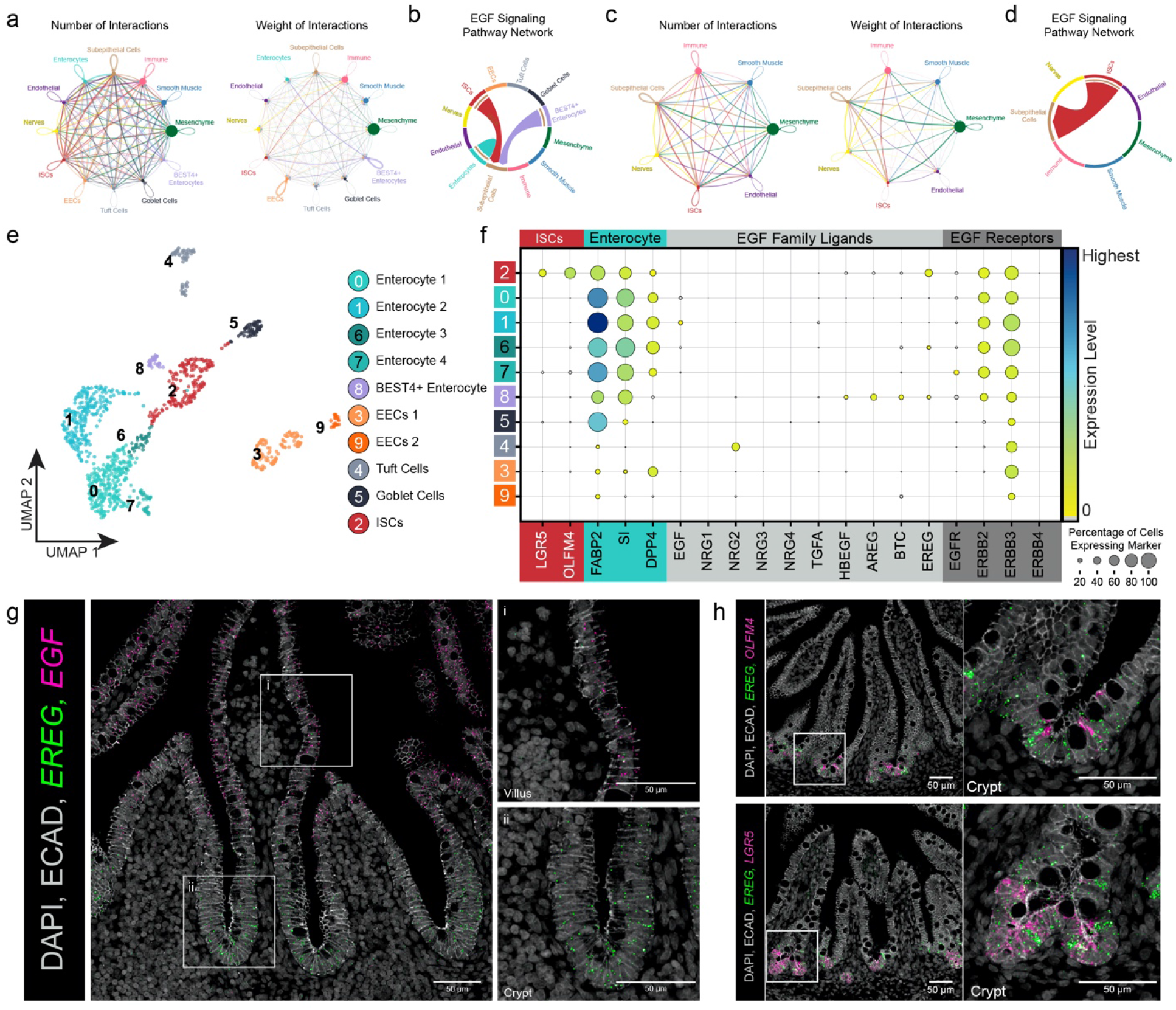
Identification of EREG as a Novel Niche Cue (a) Aggregated cell-cell communication networks showing the number of interactions (left) or total weighted interaction strengths (right) between all cell types found in 127-day and 132-day fetal tissue single cell datasets. (b) Chord diagram of predicted EGF signaling events throughout the entire dataset. (c) Aggregated cell-cell communication networks showing the number of interactions (left) or total weighted interaction strengths (right) between a subset of cell types (stem cells and all other cell types) 127-day and 132-day fetal tissue single cell datasets. (d) Chord diagram of predicted EGF signaling events throughout the subsetted dataset. (e) UMAP visualization of human fetal small intestinal epithelium (1,009 cells, n=3 biological replicates, ages 127-day and 132-day). Epithelial cluster identity was determined by expression of canonical lineage markers. (f) Dot plot visualization of stem cell markers *(LGR5, OLFM4*), enterocyte markers (*FABP2, SI, DPP4*), EGF ligands, and EGF family receptors among human fetal epithelial datasets. (g) Co-FISH/IF staining for *EREG* (green), *EGF* (pink), and ECAD (grey) in human fetal duodenum (127-day). Right of main image: Zoomed images of villus and crypt regions. (h) Co-FISH/IF staining for *EREG* (green), stem cell markers (*LGR5, OLFM4*; pink), and ECAD (grey) in human fetal duodenum (127-day).

In order to interrogate niche signaling within the stem cell compartment, we analyzed this data for receptor-ligand pairing using CellChat^35^. We initially interrogated interactions between all the cell types found in the human fetal dataset (Figure 1a) and queried specific signaling pathways known to be important for intestinal development and stem cell regulation (Figure S1a). This analysis suggested that EGF signaled between the epithelium and underlying sub-epithelial cell (SEC) compartment (Figure 1b). To examine ISC-niche interactions with more granularity, we further interrogated only stem cell interactions and observed that only one ligand was predicted to be signaling from ISCs, *EPIREGULIN* (EREG) (Figure 1c-d, Figure S1b).

CellChat analysis predicted unidirectional EGF signaling from the epithelium to the SEC; however, this prediction is not consistent with functional evidence showing that removal of EGF ligands from human or mouse organoid culture abrogates proliferation^31,36^. Therefore, we further investigated expression of all members of the EGF family of ligands and receptors in the developing stem cell niche. Consistent with prior reports, we observed expression of the receptors *EGFR* and *ErbB2* throughout the tissue (Figure S1c) and *EGF* in enterocytes (Figure 1f-g)^19^, whereas we observed robust expression of *EREG* in ISCs (Figure 1f, cluster 2 [red]). Using fluorescent *in situ* hybridization (FISH), we confirmed the spatial location of *EREG* in the crypt epithelium of the developing human intestine (Figure 1g). Co-staining with stem cell markers *LGR5* and *OLFM4* indicated that both the stem cells and surrounding epithelial cells of the crypt express *EREG* (Figure 1h). This observation suggests that EREG may have a unique role during human intestinal development.

### EREG Leads to Spatially Organized Human Epithelial Organoid Cultures

To interrogate the role of EREG in the intestinal stem cell niche, we leveraged human intestinal epithelial-only organoid cultures. We derived these cultures from isolated epithelium of developing human intestinal specimens and grew the epithelium in growth media (see Methods) supplemented with varying concentrations of EREG (100 ng/mL, 10 ng/mL, and 1 ng/mL) or matched concentrations of EGF. After 10 days, we documented the gross morphology of these structures. At all concentrations tested, EGF-grown organoids formed highly proliferative cystic structures as previously reported^37^. Conversely, EREG-grown organoids formed budded structures at lower concentrations and as protein concentration increased, structures were a mix of budded and cystic organoids (Figure 2a). To determine the EGF dose response, we sequenced 1 ng/mL EGF-grown organoids and 100 ng/mL EGF-grown organoids and found no major differences in their transcriptome or epigenome (Figure S2a). Therefore, we focused our analysis on the higher concentration since this concentration is most commonly used in the field. We quantified the topography of organoids by measuring solidity, aspect ratio, circularity, and roundness for all six conditions (Figure S2b). We observed that EREG-grown organoids had lower solidity, circularity, and roundness compared to the cystic EGF-grown organoids, quantitatively confirming the differences between budded EREG and cystic EGF-grown organoids. To measure organoid forming efficiency, we performed a quantitative single cell assay (Figure 2b) and found that EREG-grown organoids had a lower forming efficacy (1.37%) than EGF-grown organoids (1.91%), but this was not statistically significant across several biological (n=3) and technical replicates (Figure 2c).

**Figure 2.**
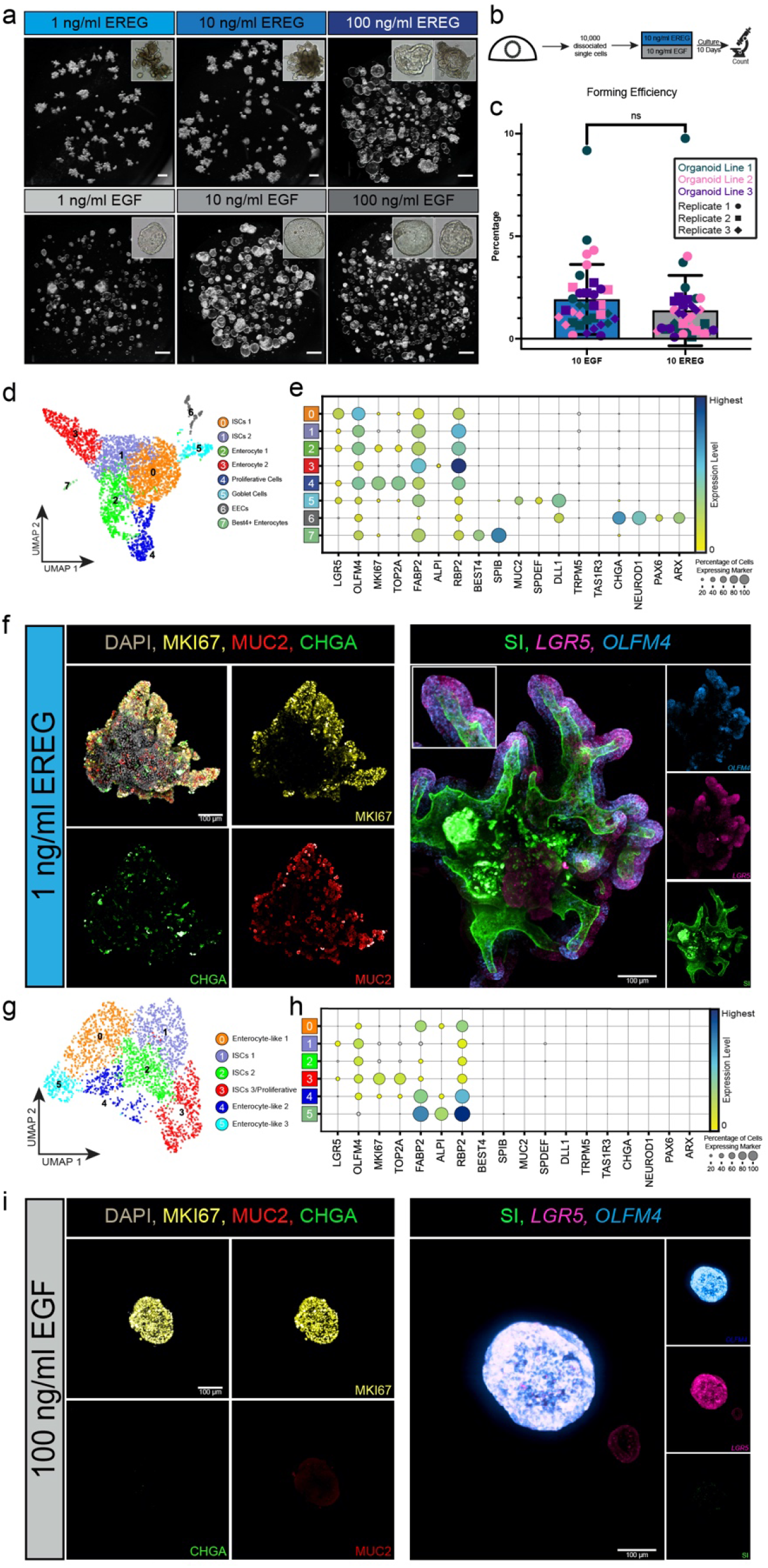
Establishment of Spatially Organized Human Epithelial Organoid Cultures (a) Brightfield images of organoids established in various concentrations of EGF or EREG. (b) Experimental design for organoid forming efficiency assay. (c) Organoid forming efficiency assay results (n=3 biological replicates with n=3 technical replicates quantified at passage 2, passage 3, and passage 4). Statistical significance was determined using a paired Welch’s two-tailed t-test using the GraphPad Prism software. (d) UMAP visualization of human fetal organoids established in EREG (1 ng/mL) at passage 1. (e) Dot plot visualization for expression of canonical markers of stem cells (*LGR5, OLFM4*), proliferative cells (*MKI67, TOP2A*), enterocytes (*FABP2, ALPI, RBP2*), BEST4+ enterocytes (*BEST4, SPIB*), goblet cells (*MUC2, SPDEF, DLL1*), tuft cells (*TRPM5, TAS1R3*), and enteroendocrine cells (*CHGA, NEUROD1, PAX6, ARX*) in EREG-grown (1ng/mL) organoids. (f) Wholemount IF staining (left) of EREG-grown (1 ng/mL) organoids for the presence of proliferation (MKI67; yellow), and differentiation into goblet cells (MUC2; red), and enteroendocrine cells (CHGA; green). Wholemount co-FISH/IF (right) for stem cell markers (*LGR5*; pink, *OLFM4*; blue) and brush border of enterocytes (sucrose isomaltose (SI); green). (g) UMAP visualization of human fetal organoids established in EGF (100 ng/ mL) organoids at passage 1. (h) Dot plot visualization for expression of canonical markers of stem cells (*LGR5, OLFM4*), proliferative cells (*MKI67, TOP2A*), enterocytes (*FABP2, ALPI, RBP2*), BEST4+ enterocytes (*BEST4, SPIB*), goblet cells (*MUC2, SPDEF, DLL1*), Tuft cells (*TRPM5, TAS1R3*), and enteroendocrine cells (*CHGA, NEUROD1, PAX6, ARX*) in EGF-grown (100 ng/mL) organoids. (i) Wholemount IF staining (left) of EGF-grown (100 ng/mL) organoids for the presence of proliferation (MKI67; yellow), and differentiation into goblet cells (MUC2; red), and enteroendocrine cells (CHGA; green). Wholemount co-FISH/IF (right) for stem cell markers (*LGR5*; pink, *OLFM4*; blue) and brush border of enterocytes (sucrose isomaltose (SI); green).

To better understand the cell composition of EREG-and EGF-grown organoids, we performed scRNA-seq on both conditions and used well-established genes to identify epithelial composition of both samples. To characterize cellular localization and spatial organization of organoids derived from both growth conditions, we also carried out 2D and 3D imaging. scRNA-seq data from EREG-grown organoids demonstrated that all cell types found in the epithelium of the developing human intestine (Figure 1) were present, including ISCs (*LGR5/OLFM4/MKI67*), goblet cells (*MUC2/SPDEF/DLL1*), enteroendocrine cells (*CHGA/NEUROD1/PAX6/ARX/REG4)*, BEST4+ enterocytes (*BEST4/SPID*) (Figure 2d-e).

However, EGF-grown organoids lacked expression of most major cell types and were instead comprised mostly of stem cells and progenitor cells beginning to differentiate into enterocyte-like cells (Figure 2g-h). Wholemount immunofluorescent staining of EREG-grown organoids demonstrated the presence of goblet and enteroendocrine cells throughout the budded structures (Figure 2f) consistent with *in vivo* localization (Figure S2c,f). Enterocytes, identified by the brush border enzyme sucrase isomaltase (SI), lined the apical wall of the central lumen.

Stem cell genes (*OLFM4, LGR5)* and proliferation marker (MKI67) were restricted to the budded regions suggesting that these structures are crypt-like domains (Figure 2f). Observations from 3D imaging were confirmed in thin sections via FISH/IF and via transmission election microscopy (Figure S2d,g). EGF-grown organoids, conversely, lacked expression of goblet, enteroendocrine and brush border markers, but had MKI67 staining throughout organoids, corresponding to ubiquitous expression of stem cell markers (*OLFM4, LGR5*) (Figure 2i and S2e). These data indicate that EREG-grown organoids recapitulate spatial organization and cellular diversity of the developing human intestine *in vitro*.

### EGF-Grown Organoids Ectopically Express Gastric Epithelial Genes

After analyzing data independently (Figure 2), we integrated scRNA-seq data from EGF and EREG-grown organoids to compare differences more directly. We observed that the two samples clustered separately (Figure 3a and Figure S3a-b), and the majority of cells occupied distinct clusters within the UMAP embedding (Figure 3b). We investigated differentially expressed genes (Table S1) and noticed higher expression of genes that are typically associated with the gastric epithelium, including TFF1 and TFF2^38,39^, in the EGF-grown organoids (Figure 3b-c and S3c), raising the possibility that EGF-grown organoids express non-intestinal genes. To interrogate this possibility, we compiled a cohort of 50 genes that are highly expressed in the developing stomach epithelium and 50 genes that are highly expressed in the developing intestinal epithelium based on a recently published Human Fetal Endoderm Atlas^33^. This allowed us to generate a ‘score’ for each sample based on stomach or intestinal gene sets. This analysis demonstrated that the EGF-grown organoids scored more highly for the stomach gene cohort, and the EREG-grown organoids scored more highly for the small intestinal gene cohort (Figure 3d-e and Table S2). To validate this finding using a complementary computational approach, we mapped EGF and EREG-grown organoids onto the entire Human Fetal Endoderm Atlas (esophagus, stomach, small and large intestine, lung, and liver), and found that a large proportion of cells from EGF-grown organoids mapped to the gastric epithelium, while EREG-grown samples primarily mapped to the small intestinal epithelium (Figure 3f). Mapping was quantified (Figure 3g) and we found that EREG-grown organoids mapped almost exclusively to the small intestine, but EGF-grown organoids had ∼40% of cells that mapped to the gastric epithelium with a small number of cells mapping to the esophagus. Immunofluorescent staining for individual stomach markers such as TFF1 (Figure 3h) and TFF2 (Figure S3d) confirmed ectopic stomach expression throughout EGF-grown organoids.

**Figure 3.**
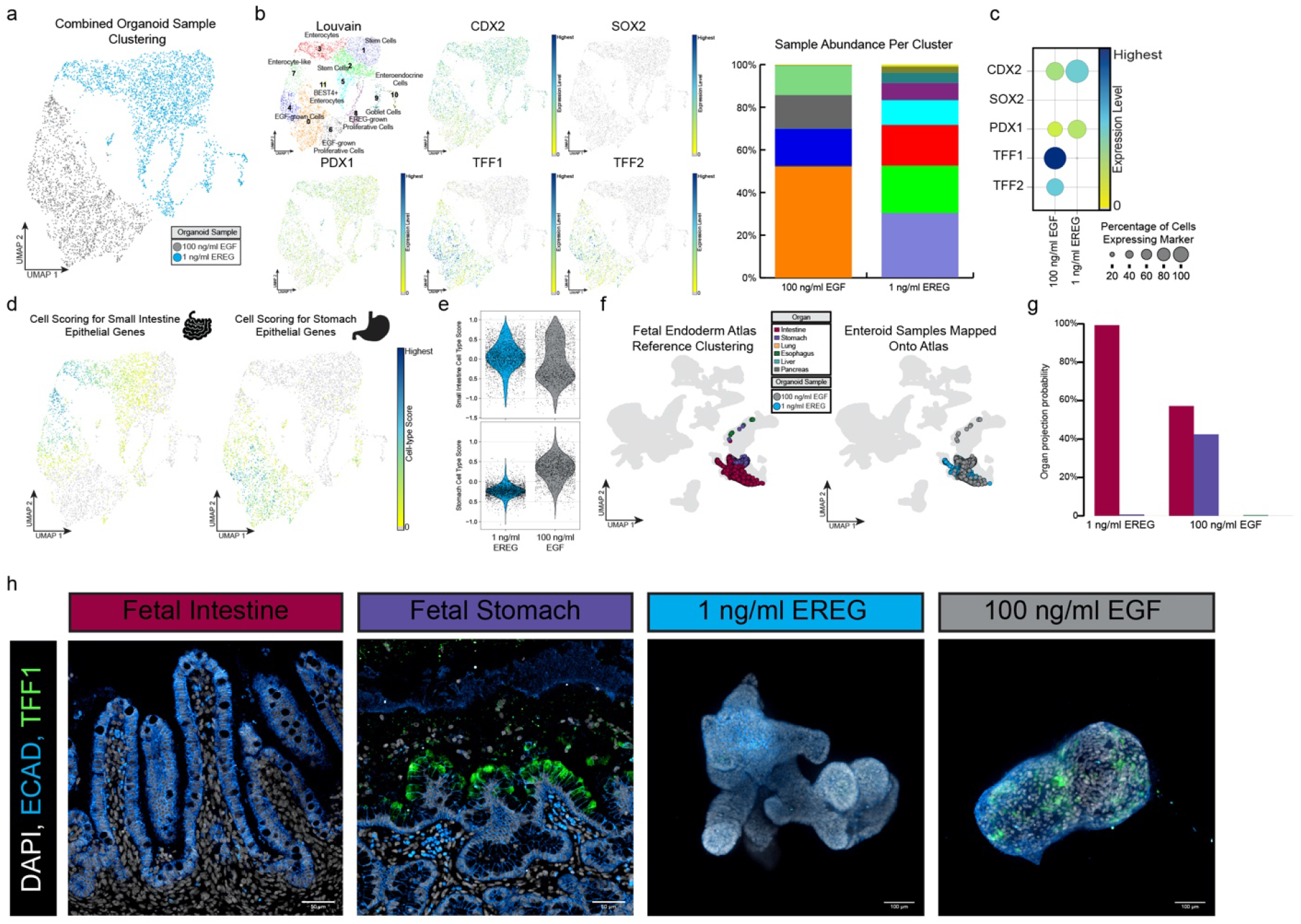
EGF-Grown Organoids Ectopically Express Gastric Epithelial Genes (a) UMAP visualization of EREG-grown (1 ng/mL) organoids (blue) and EGF-grown (100 ng/mL) organoids (grey). (b) UMAP visualization of overall Louvain clustering, and gene expression of *CDX2, SOX2, PDX1, TFF1*, and *TFF2*. Bar plot quantification of cell type abundance per Louvain cluster in each sample, colors correspond to color of cluster. (c) Dot plot quantification of gene expression shown in B, grouped by culture condition. (d) UMAP visualization of cell scoring analysis for small intestine genes (left) and stomach genes (right), see Table S2 for gene list. (e) Violin plot quantification of cell type score as shown in E (f) UMAP visualization of reference fetal atlas organ tissues^18^ (left) and organoid samples embedding onto reference map (right), blue dots represent 1 ng/mL EREG-grown cells and grey dots represent 100 ng/mL EGF-grown cells. (g) Quantification of (g). Bars represent the percentage of each sample that map to a specific tissue type. (h) Immunofluorescence staining for stomach gene TFF1 in human fetal intestine (127-day) and stomach (132-day), 1 ng/mL EREG-grown organoid (passage 1 day 10), and 100 ng/mL EGF-grown organoid (passage 1 day 10).

### EGF-Grown Organoids Have Reduced CDX2 Expression and an Altered Chromatin Landscape, Corresponding to Ectopic Stomach Gene Expression

To further understand how EGF- or EREG-culture conditions affect the intestinal stem cells, we isolated fresh crypts from fetal tissue, immediately snap froze a portion of the sample, and cultured the rest of the sample in either EREG or EGF for one passage. We then isolated nuclei and subjected all three samples to 10X multiomic sequencing to observe changes in gene expression and chromatin landscape among the samples, and to directly compare culture conditions with those of the native crypt. Data were annotated (Figure S4a-b) and epithelial cells were computationally extracted for further interrogation. A weighed nearest neighbors (WNN) graph, which represents a weighted combination of RNA and ATAC-seq modalities, was used to cluster cells^40^, and visualization was carried out via UMAP (Figure 4a). We examined the chromatin accessibility for canonical markers of different epithelial cell types, including *LGR5, CHGA, DPP4, MUC2*, and observed few differences between conditions. OLFM4 peaks were more abundant in both *in vitro* samples compared to *in vivo* (Figure S4c). Since we observed ectopic stomach gene expression in the EGF-organoids (Figure 3), we interrogated stomach gene loci in the scATAC-seq data. As predicted by expression data, we found multiple stomach loci with enhanced accessibility in EGF-grown organoids compared to both *in vivo* crypts and EREG-grown organoids (Figure S4d). Next, we interrogated the multiomic data to identify transcription factors that had differential mRNA expression as well as differentially enriched motifs within the chromatin accessibility data. This analysis revealed that several genes known to be critical for intestinal development, including *CDX2, GATA4* and *HNF4A*^41,42^, and genes with these regulatory motifs were downregulated in EGF-grown organoids when compared to fresh crypts and EREG-grown organoids (Figure 4b-c). Interestingly, while *CDX2* expression and CDX2-associated motifs were reduced in EGF-grown organoids, the individual *CDX2* gene locus accessibility was not different between groups, suggesting that reduced *CDX2* expression is not a consequence of chromatin re-organization (Figure 4d). To further interrogate global chromatin accessibility changes between groups, we treated snATAC-seq data as pseudo-bulk, and applied k-means clustering to interrogate patterns of accessibility between samples (Figure S4e). We found that EREG-grown organoids maintained many regions of open chromatin similar to *in vivo* crypts, and these regions were also closed in EGF-grown organoids (i.e. Clusters 3 and 5). Of note, when we performed motif analysis, we observed that Cluster 3 and Cluster 5 were enriched for well-established intestinal transcription factor motifs, including CDX2^41^, GATA4 and HNF4A^42^ (Figure S4f and Table S3). We also observed a large group of genomic loci (Cluster 4) with a gain-of-accessibility in the EGF-grown organoids. Motif analysis revealed FRA1 motifs, which is a transcription factor associated with gastric cancer^43^, as well as SOX2^44^, which is expressed in the early foregut. Chromatin accessibility for Cluster 4 remained low in EREG-grown organoids, although levels were slightly elevated relative to *in vivo* crypts (Figure S4e). Immunofluorescent staining of EGF- and EREG-grown organoids also suggested that EREG-grown organoids had a higher level of CDX2 protein expression as compared to EGF-grown organoids (Figure 4e). Taken together, these data suggest that EGF leads to increased accessibility at stomach gene loci, and while the chromatin accessibility of important intestinal transcription factors such as CDX2 does not change, CDX2 expression levels were reduced, corresponding to a large decrease in accessibility of regulatory motifs associated with this transcription factor.

**Figure 4.**
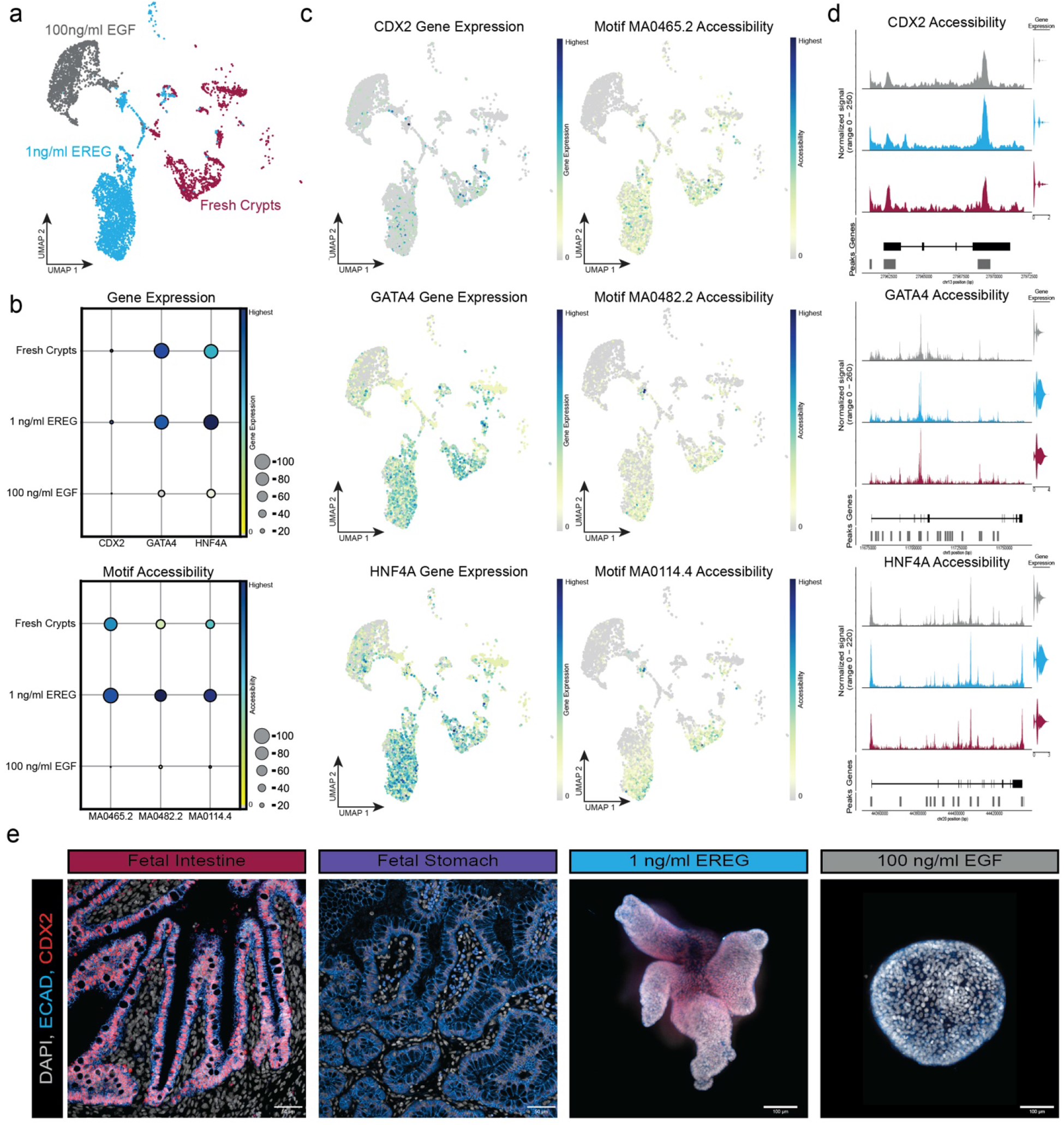
Multiomic Analysis of EGF- and EREG-grown Organoids Suggested Altered Chromatin Landscape (a) Weighed nearest neighbors (representing joint RNA and ATAC data) UMAP visualization colored by sample (fresh crypts; red, 1 ng/mL EREG (passage 0 day 10); blue, 100 ng/mL EGF (passage 0 day 10); grey). (b) Dot plot quantification of *CDX2, GATA4*, and *HNF4A* gene expression (top) and chromatin accessibility (bottom) of respective motifs for CDX2 (MA0465.2), GATA4 (MA0482.2), and HNF4A (MA0114.4). (c) UMAP visualization of *CDX2, GATA4*, and *HNF4A* gene expression (left column) and corresponding motif accessibility (right column). (d) Chromatin accessibility at CDX2, GATA4, and HNF4A loci. Diagram of each gene is along the bottom of the peaks with exons represented as black boxes and introns as lines between the exons. (e) Immunofluorescent staining for CDX2 in fetal intestine (127-day), fetal stomach (132-day), 1 ng/mL EREG-grown organoid (passage 1 day 10), and 100 ng/mL EGF-grown organoid (passage 1 day 10).

## Discussion

Here, we report EREG as a novel stem cell niche cue unique to humans and temporally controlled during development. We leveraged this cue to create a spatially organized human intestinal epithelial-only organoid system. EREG also led to gene expression and chromatin accessibility signature that remained more faithful to the native crypt epithelium compared to EGF. We also observed that the widely used EGF culture system causes ectopic stomach gene expression. The shift in chromatin landscape caused by EGF *in vitro* corresponds to a decrease in *CDX2* expression in these organoids and is also correlated with reduced chromatin accessibility of genes with CDX2 motifs. These observations are consistent with prior studies examining this master intestinal transcription factor, which causes a complete loss of intestinal identity when genetically deleted in the early murine intestine^6^, or leads to the ectopic expression of stomach genes when deleted from the intestine at later stages of development or in the adult^41^. In sum, EREG leads to an improved physiologic niche *in vitro*, giving rise to a spatially organized human epithelial organoid with crypt-like and villus-like domains, along with all major differentiated cell types of the human intestine.

## Methods

### Microscopy

All fluorescent images were taken on either a Nikon A1 confocal microscope for 2D sections, an Olympus IX83 fluorescence microscope for brightfield images, or a Nikon X1 Yokogawa Spinning Disk/Yokagawa CV8000 microscope for wholemount images. TEM sample grids were imaged on a JEOL JEM 1400 PLUS TEM. Acquisition parameters were kept consistent within the same experiment and all post-image processing was performed equally across all images of the same experiment. Images were assembled in Adobe Photoshop CC 2022.

### Tissue Processing, Staining, and Quantification

#### Tissue Processing

After harvest, tissue was immediately fixed in 10% Neutral Buffered Formalin (NBF) for 24 hours at RT on a rocker. Tissue was then washed three times in UltraPure DNase/RNase-Free Distilled Water (Thermo Fisher, Cat #10977015) for 30-60 minutes per wash depending on sample size. Next, tissue was dehydrated through an alcohol series diluted in UltraPure DNase/RNase-Free Distilled Water for 30-60 minutes per solution: 25% MeOH, 50% MeOH, 75% MeOH, 100% MeOH. Tissue was either immediately processed into paraffin blocks or stored long term at 4°C. Immediately before paraffin processing, dehydrated tissue was placed in 100% EtOH, followed by 70% EtOH, and paraffin processed in an automated tissue processor (Leica ASP300) with 1 hour solution changes overnight. Prior to sectioning, the microtome and slides were sprayed with RNase Away (Thermo Fisher, Cat#700511). 5 μm-thick sections were cut from paraffin blocks onto charged glass slides. Slides were baked for 1 hour in a 60°C dry oven (within 24 hours of performing FISH or within a week for IF). Slides were stored at RT in a slide box containing a silica desiccator packet and the seams were sealed with parafilm.

#### Immunofluorescence (IF) Protein Staining on Paraffin Sections

Tissue slides were rehydrated in Histo-Clear II (National Diagnostics, Cat#HS-202) 2x for 5 minutes each, followed by serial rinses through the following solutions 2x for 2 minutes each: 100% EtOH, 95% EtOH, 70% EtOH, 30% EtOH, and finally in double-distilled water (ddH2O) 2x for 5 minutes each. Antigen retrieval was performed using 1X Sodium Citrate Buffer diluted from a 10X stock (10X Stock: 100mM trisodium citrate (Sigma, Cat#S1804), 0.5% Tween 20 (Thermo Fisher, Cat#BP337), pH 6.0), steaming the slides for 20 minutes then washing 2x for 5 minutes in ddH2O. Slides were incubated in a humidity chamber at RT for 1 hour with blocking solution (5% normal donkey serum (Sigma, Cat#D9663) in PBS with 0.1% Tween 20). Slides were then incubated in primary antibody diluted in blocking solution at 4°C overnight in a humidity chamber. The next day, slides were washed 3x in 1X PBS for 5 minutes each and then incubated with secondary antibody with DAPI (1ug/mL) diluted in blocking solution for 1 hour at RT in a humidity chamber. Slides were then washed 3x in 1X PBS for 5 minutes each and mounted with ProLong Gold (Thermo Fisher, Cat#P369300) and imaged within a week. Stained slides were stored in the dark at 4°C. Secondary antibodies, raised in donkey, were purchased from Jackson Immuno and used at a dilution of 1:500.

#### Fluorescence in situ Hybridization (FISH) on Paraffin Sections

The FISH protocol was performed according to the manufacturer’s instructions (ACDbio, RNAscope multiplex fluorescent manual) with a 30-minute protease treatment and 15-minute antigen retrieval. For IF protein co-stains, the DAPI counterstaining final step of the FISH protocol was skipped and instead, the slides were washed 3x in 1X PBS for 5 minutes each, followed by the blocking portion of the IF protocol above.

#### Whole Mount Immunofluorescence with Antibody Staining

All tips and tubes were coated with 1% BSA in 1X PBS to prevent tissue loss. Organoids were dislodged from Matrigel using a blunt tip P1000 and transferred to a 1.5mL Eppendorf tube. 500μL of Cell Recovery Solution (Corning, Cat#354253) was added to the tube and placed on a rocker at 4°C for 50 minutes to completely dissolve Matrigel. Tube was spun at 100g for 5 minutes, after which the solution and remaining Matrigel was removed. Tissue was then fixed in 10% NBF overnight at RT on a rocker. The next day, tissue was washed 3x for 2 hours with 1mL of Organoid Wash Buffer (OWB) (0.1% Triton, 0.2% BSA in 1X PBS) at RT on a rocker.

Wash times vary (30 minutes – 2 hours) depending on tissue size. 1mL CUBIC-L (TCI Chemicals Cat. No. T3740) was added to the tube and placed on a rocker for 24 hours at 37°C. Tissue was then permeabilized for 24 hours at 4°C on rocker with 1mL Permeabilization Solution (5% Normal Donkey Serum, 0.5% Triton in 1X PBS). After 24 hours, Permeabilization Solution was removed and 500μL primary antibody (diluted in OWB) was added overnight at 4°C on a rocker. The next day, tissue was washed 3x with 1mL of OWB, 2 hours each at RT. 500μL of secondary antibody (diluted in OWB at a 1:500 dilution) was added and incubated overnight at 4°C, wrapped in foil. Tissue was washed again 3x with 1mL OWB at RT, first wash for 2 hours with an added DAPI dilution of 1:1000, then 30 minutes for the remaining washes. Samples were transferred to a 96 well glass bottom imaging plate (ThermoFisher Cat. No. 12-566-70) and then cleared and mounted with 50μL CUBIC-R (just enough to cover tissue) (TCI Chemicals Cat. No. T3741). Whole mount images were taken on a Nikon X1 Yokogawa Spinning Disk Microscope or the Yokagawa CV8000 microscope.

#### Whole Mount Fluorescent in situ Hybridization with Antibody Staining

All tips and tubes were coated with 1% BSA in 1X PBS to prevent tissue loss. Organoids were dislodged from Matrigel using a blunt tip P1000 and transferred to a 1.5mL Eppendorf tube. 500μL of Cell Recovery Solution (Corning Cat. No. 354253) was added to the tube and placed on a rocker at 4°C for 50 minutes to completely dissolve Matrigel. Tube was spun at 100g for 5 minutes, after which the solution and remaining Matrigel was removed. Tissue was fixed in 10% NBF overnight at RT on a rocker. Tissue was washed 3x for 30 minutes at RT with RNAse free water and then dehydrated using a methanol series of 25% MeOH in PBS with 1% Tween (PBST), 50% MeOH in PBST, 75% MeOH in PBST for 30 minutes to an hour depending on size. Organoids were stored in 100% MeOH at -20°C for up to 6 months. Organoids were then rehydrated using the same methanol gradient and washed in 1X PBS for the final wash.

Organoids were moved to an imaging plate for the remainder of the protocol. Using the RNAscope Multiplex Fluorescent Reagent Kit v2 (ACDbio, Cat. #323100), and leaving the PBS in the well, 3 drops of H2O2 were added at RT for 10 minutes on a rocker. Supernatant was aspirated and sample was washed 3x with ddH2O for 5 minutes each wash. 100uL of diluted RNAscope protease III (1:15 in PBS) was added for 10 minutes at 40°C in a humidity chamber. Protease was removed and sample was washed 3x in 1X PBS for 5 minutes each. 75uL probe was added and incubated for 2 hours at 40°C in the humidity chamber. Samples were washed 2x for 5 minutes in RNAscope wash buffer. From here, the remainder of the protocol was performed according to the manufacturer’s instructions (ACDbio; RNAscope Multiplex Fluorescent Manual Protocol, 323100-USM). To add an IF protein co-stain, tissue then was washed quickly with 3x PBST at RT on a rocker, wrapped in aluminum foil. Tissue was blocked with 5% NDS in 0.1% PBST for 1 hour at RT on a rocker. 500μL primary antibody (diluted in Organoid Wash Buffer (OWB) (0.1% Triton, 0.2% BSA in 1X PBS)) was added overnight at 4°C on a rocker. The next day, tissue was washed 3x with 1mL of OWB, 2 hours each at RT. Wash times vary (30 minutes – 2 hours) depending on tissue size. 500μL of secondary antibody (diluted in OWB at a 1:500 dilution) was added and incubated overnight at 4°C. Tissue was washed again 3x with 1mL OWB at RT, first wash for 2 hours with an added DAPI dilution of 1:1000, then 30 minutes for the remaining washes. Samples were transferred to a 96 well glass bottom imaging plate (ThermoFisher Cat. No. 12-566-70) and then cleared and mounted with 50μL CUBIC-R (just enough to cover tissue) (TCI Chemicals Cat. No. T3741). Whole mount images were taken on a Nikon X1 Yokogawa Spinning Disk Microscope or the Yokagawa CV8000 microscope.

#### Transmission Electron Microscopy (TEM)

EREG-grown organoids were cultured for 1 passage and small sections of whole intestine from 89-day post conception human fetal tissue were collected for TEM and prepared using conventional TEM sample preparation methods described by the University of Michigan BRCF Microscopy and Image Analysis Laboratory. Samples were fixed in 3% glutaraldehyde + 3% paraformaldehyde in 0.1M cacodylate buffer (CB), pH 7.2. Samples were washed 3 times for 15 minutes in 0.1M CB then were processed for 1 hour on ice in a post-fixation solution of 1.5% K4Fe(CN)6 + 2% OsO4 in 0.1M CB. Samples were then washed 3 times in 0.1M CB, and 3 times in 0.1M Na2 + Acetate Buffer, pH 5.2, followed by en bloc staining for 1 hour in 2% Uranyl Acetate + 0.1M Na2 + Acetate Buffer, pH 5.2. Samples were then processed overnight in an automated tissue processor, including dehydration from H2O through 30%, 50%, 70%, 80%, 90%, 95%, 100% ethanol, followed by 100% acetone. Samples were infiltrated with Spurr’s resin at a ratio of acetone: Spurr’s resin of 2:1 for 1 hour, 1:1 for 2 hours, 1:2 for 16 hours, and absolute Spurr’s resin for 24 hours. After embedding and polymerization, samples were sectioned on an ultramicrotome.

#### Organoid Shape Quantifications

To measure organoid area, solidity, aspect ratio, circularity, and roundness, brightfield images were outlined manually using the freehand selection tool in ImageJ. For each condition, six different organoids were measured three times and these measurements were graphed in Figure S2B.

#### Isolating, Establishing and Maintaining Human Fetal Epithelial Organoids

Fresh human fetal epithelium was isolated and maintained exactly as previously described^45^. Briefly, duodenal tissue was cut into smaller pieces (∼0.5-1 cm) and cut open longitudinally to expose the villi. To separate the epithelium, specimens then were incubated in dispase (StemCell Technologies, Cat#07923) for 30 minutes on ice in a Petri dish. Dispase then was removed and replaced with 100% fetal bovine serum (FBS) for 15 minutes on ice. To mechanically separate the tissue layers, 3 mLs of Advanced DMEM/F12 (Gibco, Cat#12634-028) equal to the initial volume of FBS was added to the biopsy tissue before vigorously pipetting the mixture several times. Epithelial fragments then settled to the bottom of the Petri dish where they were collected manually on a stereoscope by pipette. The epithelium was then washed with ice-cold Advanced DMEM/F12 and allowed to settle to the bottom of a 1.5 mL tube. The supernatant was withdrawn from the loose tissue pellet and replaced with Matrigel (Corning, Cat# 354234). The Matrigel containing the isolated epithelium then was gently mixed before being pipetted into individual 50 μL droplets in a 24-well plate. The plate containing the droplets then was incubated at 37°C for at least 10 minutes to allow the Matrigel to solidify before adding media (See Media Composition section below).

Once organoids were established, Matrigel droplets were dislodged from the culture plate and were pipetted into a Petri dish. Healthy organoids were manually selected from the Petri dish under a stereoscope to be bulk-passaged through a 30G needle and embedded in Matrigel (Corning, Cat#354234). For single-cell passaging, healthy organoids were manually selected under a stereoscope and dissociated with TrypLE Express (Gibco, Cat#12605-010) at 37°C for 10 minutes before filtering through 40 mm cell strainers. Cells were then counted using a hemocytometer and 10,000 cells were embedded per Matrigel droplet. After 10 days, organoid forming efficiency was calculated by taking the number of organoids that had formed and dividing it by total number of cells seeded (10,000 cells).

#### Media Composition

Culture media consisted of LWRN conditioned media generated as previously described^45,46^ and combined with human basal media (Advanced DMEM/F12 (Gibco, Cat#12634-028); Glutamax 4 mM (Gibco, Cat#35050-061); HEPES 20 mM (Gibco, Cat#15630-080); N2 Supplement (2X) (Gibco, Cat#17502-048), B27 Supplement (2X) (Fisher, Cat#17504-044), Penicillin-Streptomycin (2X) (Gibco, Cat#15140-122), N-acetylcysteine (2 mM) (Sigma, Cat#A9165-25G), Nicotinamide (20 mM) (Sigma, Cat#N0636-061)). Complete media was comprised of 25% LWRN and 75% human basal media to which rhEGF (R&D, Cat#236-EG, 100 ng/mL, 10 ng/mL, or 1 ng/mL) or rhEREG (R&D, Cat#5898-NR-050, 100 ng/mL, 10 ng/mL, or 1 ng/mL) was added.

#### Human Subjects

Normal, de-identified human fetal intestinal tissue was obtained from the University of Washington Laboratory of Developmental Biology. All human tissue used in this work was deidentified and was conducted with approval from the University of Michigan IRB.

#### Experimental Design of Organoids Cultures

The organoid experiments were designed and conducted to reduce batch effects in scRNA-seq data and dual snRNA-seq/ATAC-seq data. All experiments comparing different treatment groups (i.e., EGF or EREG) were carried out in parallel, with experiments and treatments being conducted at the same time. Cells were harvested and dissociated into single cell suspensions or single nuclei suspensions in parallel (more details below). Since the 10X Chromium system allows parallel processing of multiple samples at a time, cells were captured (Gel bead-in-Emulsion - GEMS) and processed (i.e., library prep) in parallel by the University of Michigan Advanced Genomics Core. All samples were sequenced across the same lane(s) on a Novaseq 6000 for both transcriptional and epigenomic data.

### Single Cell Experiments

#### Single Cell Dissociation

To dissociate organoids to single cells, matched healthy organoids for the same human sample in different media conditions were manually collected under a stereoscope. Following collection, dissociation enzymes and reagents from the Neural Tissue Dissociation Kit (Miltenyi, 130-092-628) were used, and all incubation steps were carried out in a refrigerated centrifuge pre-chilled to 10°C unless otherwise stated. All tubes and pipette tips used to handle cell suspensions were pre-washed with 1% BSA in 1X HBSS to prevent adhesion of cells to the plastic. Tissue was treated for 15 minutes at 10°C with Mix 1 and then incubated for 10-minute increments at 10°C with Mix 2, and interrupted by agitation through pipetting with a P1000 pipette until fully dissociated. Cells were filtered through a 70 μm filter coated with 1% BSA in 1X HBSS, spun down at 500g for 5 minutes at 10°C and resuspended in 500mL 1X HBSS (with Mg2+, Ca2+).

Cells were spun down (500g for 5 minutes at 10°C) and washed twice by suspension in 2 mL of HBSS + 1% BSA, followed by centrifugation. Cells were counted using a hemocytometer, then spun down and resuspended to reach a concentration of 1,000 cells/μL and kept on ice. Single cell libraries were immediately prepared on the 10x Chromium by the University of Michigan Advanced Genomics Core facility with a target capture of 5,000 cells. A full, detailed protocol of tissue dissociation for single cell RNA sequencing can be found at http://www.jasonspencelab.com/protocols.

#### Single Cell Library Preparation and Transcriptome Alignment

All scRNA-seq sample libraries were prepared with the 10x Chromium Controller using v3 chemistry (10x Genomics, Cat# 1000268). Sequencing was performed on a NovaSeq 6000 with targeted depth of 100,000 reads per cell. Default alignment parameters were used to align reads to the pre-prepared human reference genome (hg19) provided by the 10X Cell Ranger pipeline. Initial cell demultiplexing and gene quantification were also performed using the default 10x Cell Ranger pipeline.

#### Single Cell Data Analysis

All scRNA-seq analysis downstream of gene quantification were done using Scanpy^47^ with the 10x Cell Ranger derived gene by cell matrices. For primary human tissue sample analysis in Figure 1, 127-day and 132-day post conception duodenum (3 samples) and ileum (1 sample) were used. All samples were filtered to remove cells with less than 800 or greater than 3,500 genes, or greater than 12,000 unique molecular identifier (UMI) counts per cell. De-noised data matrix read counts per gene were log normalized prior to analysis. After log normalization, highly variable genes were identified and extracted, and batch correction was performed using the BBKNN algorithm. The normalized expression levels then underwent linear regression to remove the effects of total reads per cell and cell cycle genes, followed by a z-transformation. Dimension reduction was performed using principal component analysis (PCA) and then uniform manifold approximation and projection (UMAP) on the top 11 principal components (PCs) and 15 nearest neighbors for visualization on 2 dimensions. Clusters of cells within the data were calculated using the Louvain algorithm within Scanpy with a resolution of 0.3.

Epithelial cells were identified using canonically expressed genes and extracted from a data matrix to include 1,009 intestinal epithelial cells from all samples. The extracted epithelial cell matrix again underwent log normalization, variable gene extraction, z-transformation and dimension reduction to be displayed in the UMAP seen in Figure 1 (ArrayExpress: in progress).

For Figure 2, the EREG-grown samples were filtered to remove cells with less than 1,200 or greater than 8,500 genes, or greater than 65,000 unique molecular identifier (UMI) counts per cell, and 0.1 mitochondrial cell counts. Data matrix read counts per gene were log normalized prior to analysis. After log normalization, highly variable genes were identified and extracted. Data was then scaled by z-transformation. Dimension reduction was performed using principal component analysis (PCA) and then uniform manifold approximation and projection (UMAP) on the top 10 principal components (PCs) and 15 nearest neighbors for visualization. Clusters of cells within the data were calculated using the Louvain algorithm within Scanpy with a resolution of 0.5. The EGF-grown samples were filtered to remove cells with less than 1,250 or greater than 8,500 genes, or greater than 65,000 unique molecular identifier (UMI) counts per cell, and 0.1 mitochondrial cell counts. Data matrix read counts per gene were log normalized prior to analysis. After log normalization, highly variable genes were identified and extracted. Data was then scaled by z-transformation. Dimension reduction was performed using principal component analysis (PCA) and then uniform manifold approximation and projection (UMAP) on the top 10 principal components (PCs) and 15 nearest neighbors for visualization. Clusters of cells within the data were calculated using the Louvain algorithm within Scanpy with a resolution of 0.4.

For Figure 3, all organoid samples were filtered to remove cells with less than 1,200 or greater than 7,500 genes, or greater than 50,000 unique molecular identifier (UMI) counts per cell, and 0.1 mitochondrial cell counts. Data matrix read counts per gene were log normalized prior to analysis. After log normalization, highly variable genes were identified and extracted. Data was then scaled by z-transformation. Dimension reduction was performed using principal component analysis (PCA) and then uniform manifold approximation and projection (UMAP) on the top 12 principal components (PCs) and 15 nearest neighbors for visualization. Clusters of cells within the data were calculated using the Louvain algorithm within Scanpy with a resolution of 0.3.

#### Cell Scoring Analysis

Cells were scored based on expression of a set of 50 marker genes per epithelial tissue type. Gene lists were compiled based on the previously published top 50 most differentially expressed genes from *in vivo* human epithelial cells of the small intestine and stomach^33^. See Table S2 for gene lists. After obtaining the log normalized and scaled expression values for the data set, scores for each cell were calculated as the average z-score within each set of selected genes.

#### Receptor-Ligand Analysis

*In silico* ligand-receptor analysis was performed using the CellChat R Package (https://github.com/sqjin/CellChat) on the 127-day and 132-day human fetal intestine datasets. Log transformed and normalized counts along with cluster identities were used as input for the analysis using standard parameters and the standard workflow to analyze a single dataset was used.

#### Endoderm Atlas

Reference map embedding to the Human Fetal Endoderm Atlas^33^ was performed using the scoreHIO R Package (https://github.com/Camp-Lab/scoreHIO). Organoid samples were processed following the preprocessing steps outlined above in the single cell data analysis section and then put through the basic workflow outlined in the package to map organoid cells onto the reference map and quantify their identity.

### Dual snRNA-seq/ATAC-seq Experiments

#### Single Nuclei Isolation and Permeabilization

Nuclei were isolated and permeabilized in accordance with 10x Genomics’ protocol for Nuclei Isolation from Complex Tissues for Single Cell Multiome ATAC + Gene Expression Sequencing. Briefly, tissue was minced into smaller fragments and then put into NP40 lysis buffer (Fisher, Cat# PI28324). Tissue was homogenized with a pellet pestle 15x by hand, then incubated in the lysis buffer for 5 minutes. The suspension was then passed through a 70μm strainer followed by a 40μm strainer. The suspension was then centrifuged at 500g for 5 minutes at 4°C. The supernatant was removed, and the pellet was washed 2x with PBS + 1% BSA and centrifuged at 500g for 5 minutes at 4°C after each wash. The pellet was then incubated in 0.1X Lysis Buffer for 2 minutes to permeabilize the sample. The sample was then centrifuged at 500g for 5 minutes at 4°C, supernatant was removed, and resuspended in diluted nuclei buffer.

#### Single Nuclei/ATAC Library Preparation and Transcriptome Alignment

All multiomic sample libraries were prepared with the Single Cell Multiome ATAC + Gene Expression v1 chemistry. Sequencing was performed on a NovaSeq 6000 with a targeted depth of 25,000 reads per cell for both ATAC and RNA. Default alignment parameters were used to align reads to the pre-prepared hg38 human reference genome provided by the 10X Cell Ranger ARC pipeline. Initial cell demultiplexing and gene quantification were also performed with the default 10x Cell Ranger ARC pipeline.

#### Single Nuclei Analysis

All multiomic analysis downstream of gene quantification were done using Seurat and the Signac extension with the 10x Cell Ranger ARC derived gene by cell matrices. For sample analysis in Figure 4 and Figure S2A, all samples were filtered to remove cells with more than 100 but less than 15,000 unique molecular identifier (UMI) counts per cell for RNA assay and more than 100 or less than 45,000 UMIs for the ATAC assay. A TSS enrichment score of 0.5 or over was also used in filtering. Next, peak calling was performed using MACS2^48^. Normalization of gene expression and dimension reduction was performed using principal component analysis (PCA). DNA accessibility assay was processed by latent semantic indexing (LSI) and a WNN graph was calculated representing a weighted combination of RNA and ATAC-seq modalities. We use this graph to perform cell type clustering and visualized it using a UMAP visualization. Epithelial cells were identified based on their expression of the pan-epithelial marker *CDH1* and lack of expression of pan-mesenchymal marker *VIM*. From here, motif analysis and peak visualization were performed.

## Author Contributions

C.J.C. and J.R.S. conceived the study. J.R.S. supervised the research. C.J.C. and O.P.C., performed computational analyses. C.J.C., J.H.W., O.P.C., Q.Y., J.G.C., M.P.V., and J.R.S. interpreted computational results and provided support. A.W., E.M.H., Y.-H.T., and M.M.C. developed tissue dissociation methods and generated single-cell RNA sequencing data. C.J.C developed tissue dissociation method for multiomic analysis and the joint sequencing data. E.M.H., C.W.S., C.J.C., C.Z., and J.Z.S. performed and analyzed all staining experiments and imaging. C.W.S., Y.-H.T., A.W., and C.J.C. performed enteroid experiments. C.J.C., C.W.S., Y.- H.T., A.W., M.M.C., M.E., and E.M.H. analyzed and interpreted enteroid experiments. R.Z., B.P., C.T., O.K., and I.G. provided critical material resources for this work. C.J.C., and E.M.H., assembled figures. C.J.C., E.M.H., and J.R.S. wrote the manuscript. C.J.C., E.M.H., C.W.S., Y.- H.T., A.W., J.H.W., M.E., and C.L., contributed methods. All authors edited, read, and approved the manuscript.

## Code Availability

Code used for computational analysis in this manuscript can he found at: https://github.com/jason-spence-lab/Childs_2022

**Supplemental Figure 1.**
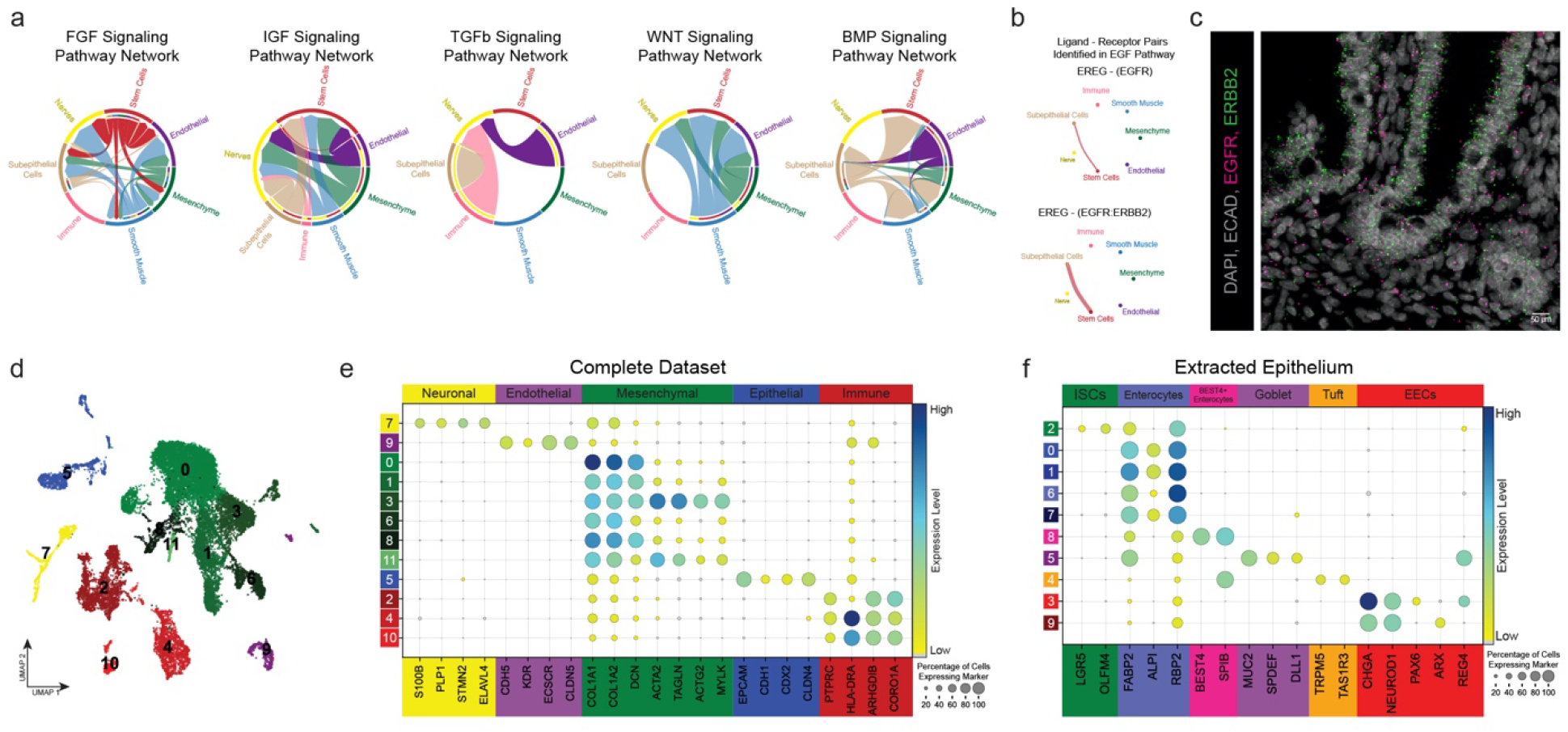
Whole Tissue Cell Cluster Annotation and Receptor-Ligand Analysis of Developing Human Small Intestine at Single Cell Resolution. Related to Figure 1. (a) Chord diagram of predicted FGF, IGF, TGFb, WNT, and BMP signaling events between intestinal stem cells and additional non-epithelial lineages. (b) Diagram showing the contribution of ligand-receptor pairing to chord diagram in Figure 1D. EREG was the only ligand predicted in the stem cell cluster. (c) Co-FISH/IF staining for *EGFR* (pink), *ErbB2* (green), and ECAD (grey) in human fetal duodenum (127-day). (d) UMAP visualization of entire merged fetal datasets colored broadly by cell class (neurons; yellow, endothelial; purple, mesenchyme; green, epithelium; blue, and immune; red) datasets include 2 biological replicates, ages 127-day (two duodenal samples) and 132-day (one duodenal and one ileum) with 18,100 cells total. (e) Dot plot of entire fetal datasets highlighting expression of canonical lineage genes that were used for cluster annotation. (f) Dot plot illustrating expression of canonical epithelial lineage markers within fetal epithelial subclusters.

**Supplemental Figure 2.**
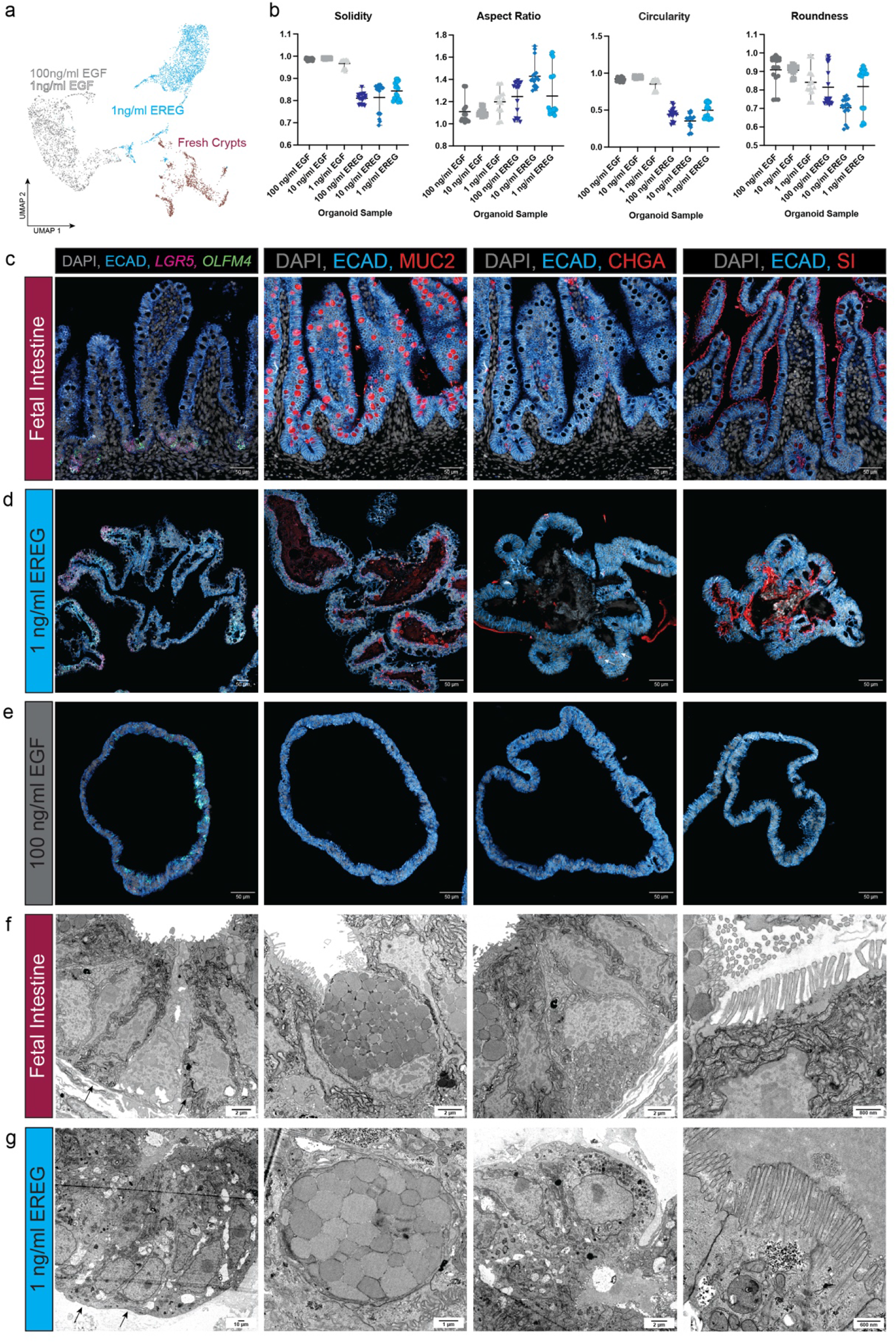
Quantification and Additional Staining of EREG and EGF-grown Organoids. Related to Figure 2. (a) Weighed nearest neighbors (representing joint RNA and ATAC data) UMAP visualization colored by sample (fresh crypts; red, 1 ng/mL EREG; blue, 1 ng/mL and 100 ng/mL EGF; grey). Overlap of EGF conditions suggests transitional and epigenomic similarity. (b) Quantification of solidity, aspect ratio, circularity, and roundness of organoids grown in varying doses of EGF or EREG. Six organoids grown for one passage to 10 days were measured three times. (c) FISH and immunofluorescent staining in fetal intestine (127-day) tissue sections for stem cells *(LGR5, OLFM4*), goblet cells (MUC2), enteroendocrine cells (CHGA), and brush border of enterocytes (SI). (d) FISH and immunofluorescent staining of 1 ng/mL EREG-grown organoids specimens for stem cells *(LGR5, OLFM4*), goblet cells (MUC2), enteroendocrine cells (CHGA), and brush border of enterocytes (SI). (e) FISH and immunofluorescent staining of 100 ng/mL EGF-grown organoids specimens for stem cells *(LGR5, OLFM4*), goblet cells (MUC2), enteroendocrine cells (CHGA), and brush border of enterocytes (SI). (f-g) TEM imaging of 1 ng/mL EREG-grown organoids (f) and human fetal intestine (89-day) (g) specimens. Intracellular characteristics were used to classify the presence of stem cells (black arrows), goblet cells, enteroendocrine cells, and brush border of enterocytes.

**Supplemental Figure 3.**
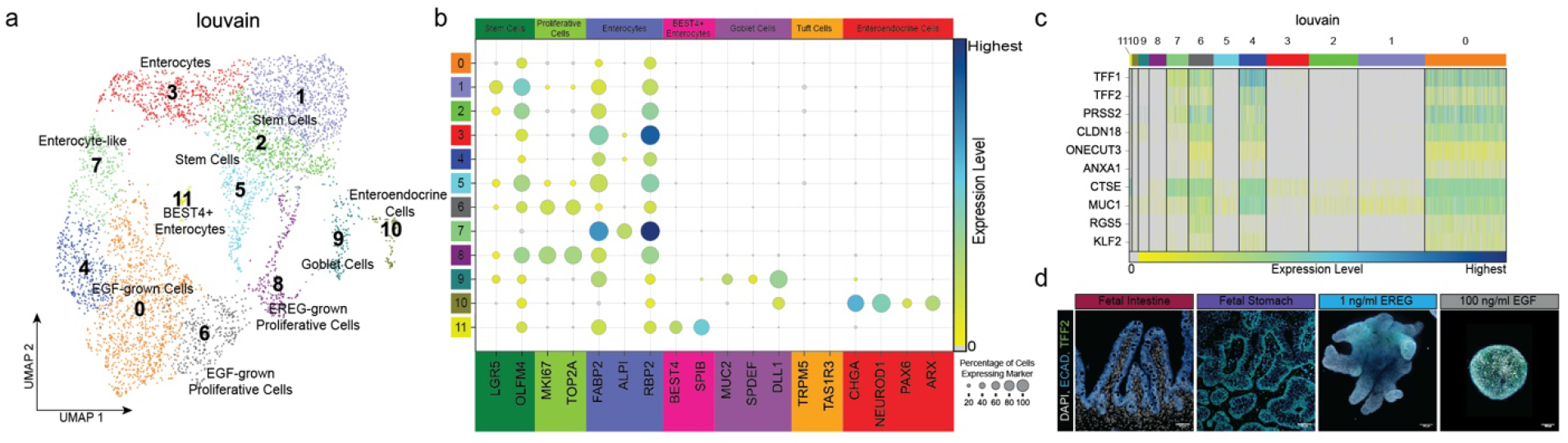
Cell Cluster Annotation of Organoid Samples and Ectopic Stomach Expression by scRNA-seq and Immunofluorescence Imaging. Related to Figure 3. (a) UMAP visualization of Louvain clustering of 1 ng/mL EREG and 100 ng/mL EGF organoids. Clusters annotated using canonically expressed marker genes when possible, otherwise growth condition is noted in cluster name (i.e. EGF-grown cells). (b) Dot plot visualization of canonically expressed marker gene used to annotated Louvain clustering in (A). These include stem cells (*LGR5, OLFM4*), proliferative cells (*MKI67, TOP2A*), enterocytes (*FABP2, ALPI, RBP2*), BEST4+ enterocytes (*BEST4, SPIB*), goblet cells (*MUC2, SPDEF, DLL1*), tuft cells (*TRPM5, TAS1R3*), and enteroendocrine (*CHGA, NEUROD1, PAX6, ARX*). (c) Heatmap showing ectopic stomach gene expression in Louvain clusters with most expression occurring in EGF-grown cell clusters (0, 4, 6, 7). (d) Immunofluorescence staining for stomach marker TFF2 (green) counterstained with ECAD (blue) and DAPI (grey) in human fetal intestine (127-day), human fetal stomach (132-day), 1 ng/mL EREG-grown organoid (passage 1 day 10), and 100 ng/mL EGF-grown organoid (passage 1 day 10).

**Supplemental Figure 4.**
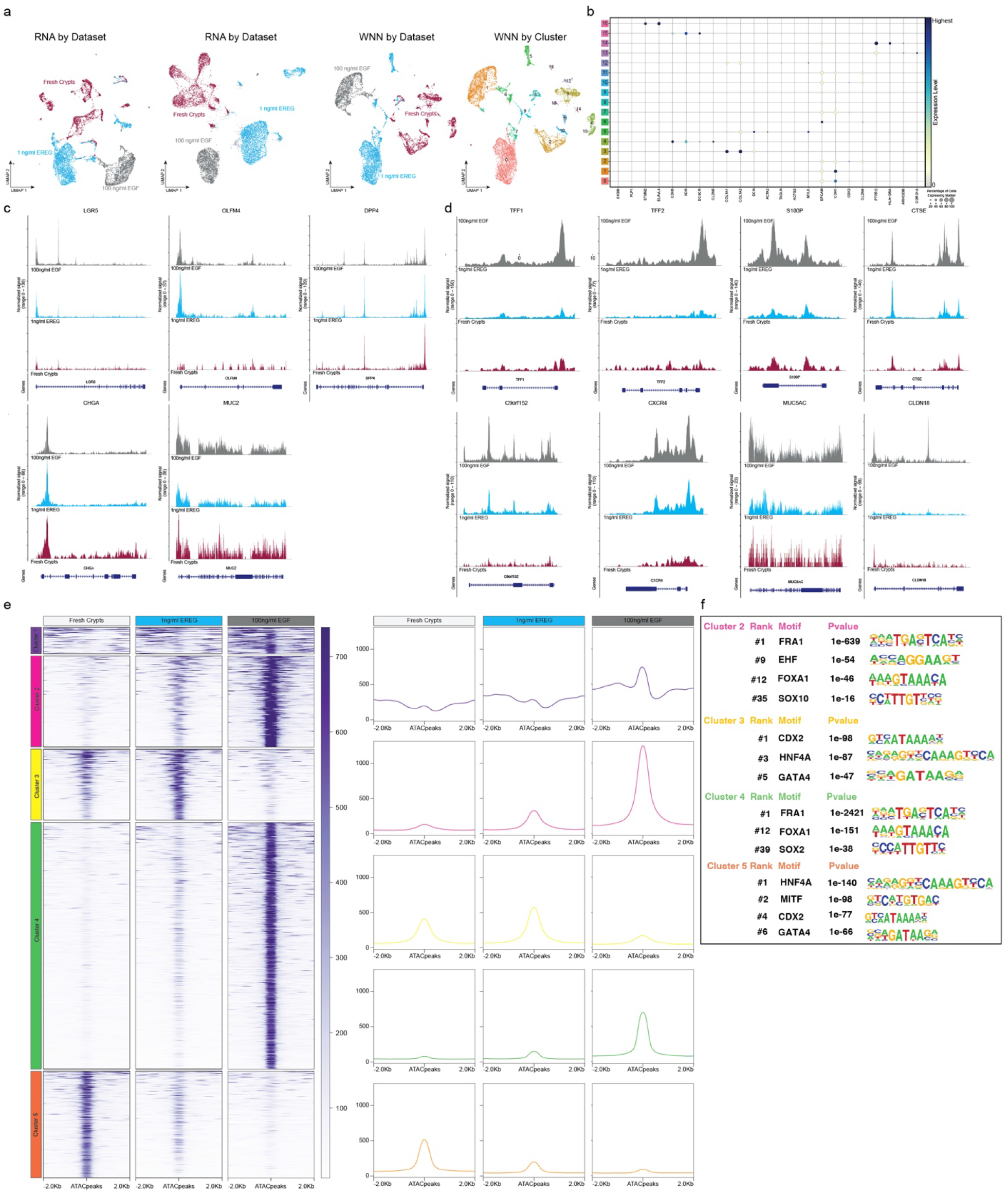
Characterization of Multiomic Analysis and k-means Clustering. Related to Figure 4. (a) UMAP visualization of colored by sample (from left to right) visualized by RNA, ATAC, WNN (joint RNA/ATAC), and WNN by clusters. (b) Dot plot of canonically expressed genes used for cluster identity in A (c) Chromatin accessibility at genes associated with small intestine epithelium (d) Chromatin accessibility at genes associated with the gastric epithelium (e) k-means clustering of peaks in control fresh crypts, 1 ng/mL EREG, and 100 ng/mL EGF. (left) Heatmap of accessibility in these regions, (right) graphical schematic summary of overall pattern of peaks shown on the right. (f) Motif analysis of clusters of interest from k-means clustering in (e) including motif and P-value. Full list of ranked motifs can be found in Table S3.

